## Supplementary figures and images for "Regional heterogeneities of oligodendrocytes underlie biased Ranvier node spacing along single axons in sound localization circuit"

### Supplemental Data 1

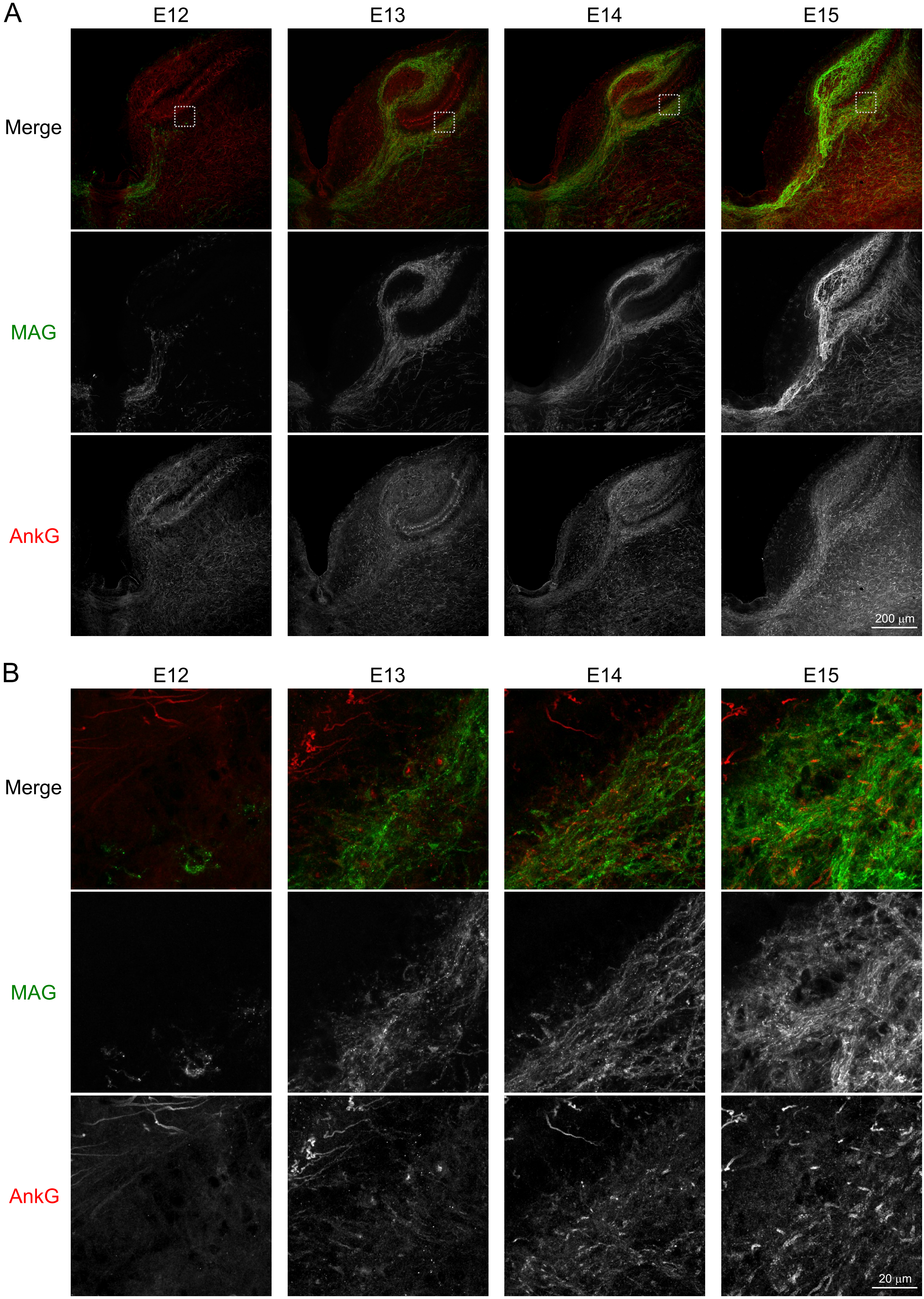


Figure 2—figure supplement 1

### Supplemental Data 2

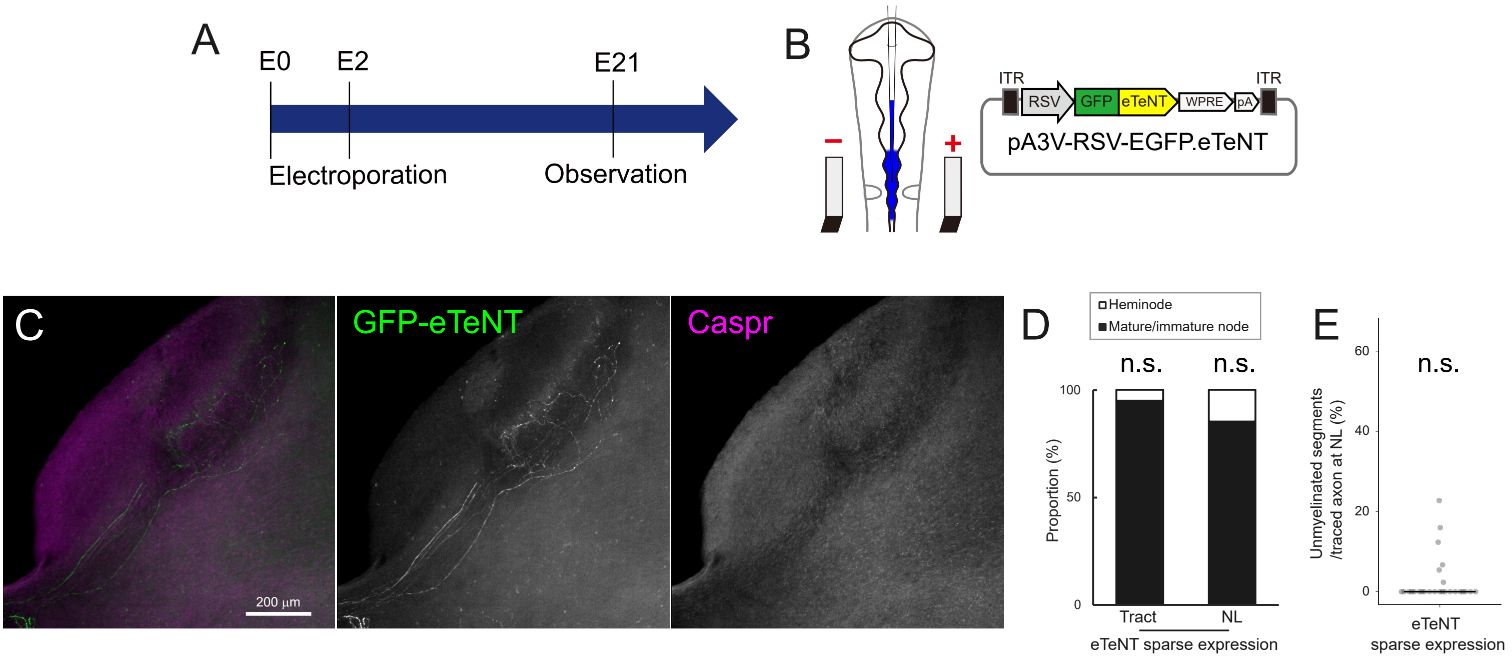


Figure 6—figure supplement 1
